# *Acinetobacter baumannii* NCIMB8209: A rare environmental strain displaying extensive insertion sequence-mediated genome remodeling resulting in the loss of exposed cell structures and defensive mechanisms

**DOI:** 10.1101/2020.04.30.071514

**Authors:** Guillermo D. Repizo, Martín Espariz, Joana L. Seravalle, Juan Ignacio Díaz Miloslavich, Bruno A. Steimbrüch, Howard A. Shuman, Alejandro M. Viale

**Affiliations:** Instituto de Biologia Molecular y Celular de Rosario (IBR, CONICET), Departamento de Microbiologia, Facultad de Ciencias Bioquimicas y Farmaceuticas, Universidad Nacional de Rosario, Rosario, Argentina; Department of Microbiology, University of Chicago, Chicago, Illinois, USA

**Keywords:** Environmental *Acinetobacter baumannii*, pre-antibiotic era *Acinetobacter baumannii*, environmental reservoirs, comparative genomics, insertion sequences, virulence factors, comparative genomics

## Abstract

*Acinetobacter baumannii* represents nowadays an important nosocomial pathogen of poorly defined reservoirs outside the clinical setting. Here we conducted whole-genome sequencing analysis of the *Acinetobacter sp*. NCIMB8209 collection strain, isolated in 1943 from the aerobic degradation (retting) of desert guayule shrubs. NCIMB8209 contained a 3.75 Mb chromosome and a plasmid of 134 kb. Phylogenetic analysis based on core genes indicated NCIMB8209 affiliation to *A. baumannii*, a result supported by the identification of a chromosomal *bla*_OXA-51_-like gene. Seven genomic islands lacking antimicrobial resistance determinants, 5 regions encompassing phage-related genes and, notably, 93 insertion sequences (IS) were found in this genome. NCIMB8209 harbors most genes linked to persistence and virulence described in contemporary *A. baumannii* clinical strains, but many of them encoding components of surface structures are interrupted by IS. Moreover, defense genetic islands against biological aggressors such as type 6 secretion systems or crispr/cas are absent from this genome. These findings correlate with a low capacity of NCIMB8209 to form biofilm and pellicle, low motility on semisolid medium, and low virulence towards *Galleria mellonella* and *Caenorhabitis elegans*. Searching for catabolic genes and concomitant metabolic assays revealed the ability of NCIMB8209 to grow on a wide range of substances produced by plants including aromatic acids and defense compounds against external aggressors. All the above features strongly suggest that NCIMB8209 has evolved specific adaptive features to a particular environmental niche. Moreover, they also revealed that the remarkable genetic plasticity identified in contemporary *A. baumannii* clinical strains represents an intrinsic characteristic of the species.

**IMPORTANCE:** *Acinetobacter baumannii* (*Ab*) is an ESKAPE opportunistic pathogen, with poorly defined natural habitats/reservoirs outside the clinical setting. *Ab* arose from the Acb complex as the result of a population bottleneck, followed by a recent population expansion from a few clinically-relevant clones endowed with an arsenal of resistance and virulence genes. Still, the identification of virulence traits and the evolutionary paths leading to a pathogenic lifestyle has remained elusive, and thus the study of non-clinical (“environmental”) *Ab* isolates is necessary. We conducted here comparative genomic and virulence studies on *Ab* NCMBI8209 isolated in 1943 from the microbiota responsible of the decomposition of guayule, and therefore well differentiated both temporally and epidemiologically from the nowadays predominant multidrug-resistant strains. Our work provides insights on the adaptive strategies used by *Ab* to escape from host defenses, and may help the adoption of measures aimed to limit its further dissemination.

## INTRODUCTION

The genus *Acinetobacter*, family *Moraxellaceae*, class *Gammaproteobacteria*, is characterized by Gram-negative aerobic coccobacilli of ubiquitous environmental distribution and large metabolic capabilities (1). Among the genus, the phylogenetically closely related species composing the *A. calcoaceticus*-*A. baumannii* (Acb) complex represent nowadays important opportunistic pathogens (2). Infections due to *A. baumannii* in particular, rarely reported in healthcare settings before the 1970s, rapidly increased in importance with the global spread of a limited number of epidemic clonal complexes (CC) possessing multidrug-resistance (MDR) phenotypes (2, 3). Strains composing the CCs generally contain a variety of genetic material obtained by horizontal gene transfer such as plasmids and chromosomally-located genomic islands (GI), including resistance islands (RI) encompassing different transposons and integrons which play pivotal roles in both antimicrobial and heavy metal resistances (2, 4-6). CC strains also carry a large repertoire of insertion sequences (ISs) capable of mediating gene inactivation as well as genome rearrangements, including deletions and inversions with strong adaptive significances (2, 6-8). The combination of the above factors, added to the intrinsic resistance of *A. baumannii* to desiccation and fever-associated temperatures (9), are considered main factors of persistence of this pathogen in the nosocomial environment (2, 7).

Whole-genome sequence (WGS) comparisons have become commonplace in examining strain-to-strain variability and in comparing pathogenic strains with environmental relatives of a given species, in efforts to identify underlying genetic determinants and mechanisms responsible for phenotypic dissimilarities. When applied to *A. baumannii*, these approaches provided valuable information on the clinical population structure of this species, its potential virulence factors, and the origins and acquisition of antimicrobial resistance determinants (2, 4, 6, 7, 10, 11). However, the identification of virulence traits both within and between CC members has remained elusive, suggesting a complex and even multifactorial nature of mechanisms involved (2, 3, 7). The importance of a deeper genomic study of non-clinical (“environmental”) isolates has therefore recently been emphasized, to clarify both the virulence potential of the aboriginal *A. baumannii* population and the evolutionary paths that led towards a pathogenic lifestyle (2). Nevertheless, the natural habitats and potential reservoirs of *A. baumannii* outside the clinical setting have remained poorly defined, and are only beginning to be elucidated (2, 11-13).

We have recently traced the origins of the collection strain *A. baumannii* DSM30011 (11) to an isolate originally classified as *Achromobacter lacticum*, which was obtained prior to 1944 from the enriched microbiota responsible of the aerobic decomposition of the resinous desert shrub guayule (14, 15). WGS of this strain and subsequent comparative analysis with other 32 complete clinical *A. baumannii* genomes revealed the presence of 12 unique accessory chromosomal regions in DSM30011, including five regions containing phage-related genes, five regions encoding toxins related to the type-6 secretion system, and one region encompassing a novel CRISPR/cas cluster (11). Expectedly from an environmental isolate obtained before the massive introduction of antimicrobial therapy in the 1950s (2), DSM30011 showed a general antimicrobial susceptibility phenotype and no RI in its genome (11). Notably, the genome of DSM30011 lacked complete IS elements, although some remnants could be predicted on it.

Remarkably still, most genes and regulatory mechanisms linked to persistence and virulence in pathogenic *Acinetobacter* species were identified in the DSM30011 genome. Moreover, several gene clusters encoding catabolic pathways related to the degradation of plant defenses were found, thus suggesting that plants may provide an effective niche for *A. baumannii* (11) as reported for other species of the genus (11). In turn, it also opens the possibility that phytophagous insects feeding on these plants (16-22) may represent both reservoirs and vectors for the dissemination of *A. baumannii* in the environment.

During the original isolation of DSM30011, two companion isolates also classified as *A. lacticum* displaying similar phenotypic characteristics were also described (14). Only one of them was saved in collections, and we obtained a replica of this isolate (see table S1 in 11) which had been deposited on the National Collection of Industrial Food and Marine Bacteria, Aberdeen, Scotland under the strain designation NCIMB8209, aiming to perform WGS comparative studies with other completely sequenced *Acinetobacter* genomes. We report here the result of these analyses as well as some phenotypic characteristics of this strain, which may provide clues into the environmental reservoirs, genomic diversity, and virulence potential of the pre-antibiotic era *A. baumannii* population.

## METHODS

### Bacterial strains and growth conditions

*Acinetobacter* sp. strain NCIMB8209 was obtained from the National Collection of Industrial Food and Marine Bacteria, Aberdeen, Scotland. For details on the original isolation and different denominations assigned to this strain by various collections see our previous publication (11) under “Tracing DSM30011 origins”.

The use of carbon or carbon-and-nitrogen sources by the *A. baumannii* strains NCIMB8209, DSM30011, or ATCC 17978 was conducted on solid BM2 minimal medium (62 mM potassium phosphate (pH 7.0), 7 mM (NH4)2SO4, 0.5 mM MgSO4, 10 μM FeSO4) supplemented with 1.5 % Difco Bacteriological Agar and 0.2 % (w/v) of the tested substrates (23). The plates were incubated at 30 °C for 48-96 h before colony growth was inspected. Other bacterial strains such as *Acinetobacter baylyi* ADP1 (24) and *Pseudomonas aeruginosa* PAO1 (25) were also included in these tests as controls for the utilization of particular compounds.

Plates containing 0.3% agarose, 10 g/L tryptone and 5 g/L NaCl were used as a tool to detect cell motility on a semisolid surface. The plates were inoculated on the surface with bacteria lifted from overnight LB agar cultures using flat-ended sterile wooden sticks or depositing 10 µl of LB cultures grown to an optical density at 600 nm (OD600) of 0.1. Plates were incubated for 30 h at 30°C in the dark before inspection and recording.

Tolerance to H_2_O_2_ was performed as previously described (26). Briefly, bacterial cultures were collected at 0.6 OD600 nm and subjected to serial dilutions. Aliquots of 10 µL were then loaded onto LB agar plates, supplemented with 400 µM H_2_O_2_, and incubated for 30 h at 30°C before inspection.

### NCIMB8209 genome sequencing and annotation

NCIMB8209 DNA was isolated using a commercial kit (Wizard Genomic DNA purification kit, Promega), following the manufacturer’s instructions. The genomic sequence was obtained using a hybrid strategy combining PacBio Sequel (MrDNALab, Shallowater, Texas, USA) and Illumina MiSeq (single-end reads, University of Chicago genomic facilities) methods. A total of 3,238,476,791 reads were generated with the PacBio Sequel approach (depth of coverage ∼834X), which were subjected to quality assessment prior to *de novo* genome assembly using Falcon (27) followed by further polishment by the Arrow algorithm. In turn, the sequence data generated by the Illumina strategy was assembled into contigs using Spades version 3.0.11 (28), and used to close the gaps left by the PacBio Sequel strategy. The replication origin (*oriC*) was predicted with OriFinder (29). The finally assembled NCIMB8209 chromosome was annotated using the pipeline available at NCBI and deposited at DDBJ/ENA/GenBank under the accession CP028138. In turn, the plasmid sequences derived from the above sequencing data were deposited in the GenBank nucleotide sequence database under accession number CP028139.

The presence of IS elements in the NCIMB8209 genome was determined using IS Finder (30) (https://www-is.biotoul.fr/). Novel ISs were deposited in the ISSaga database (31). Putative prophage sequences were identified by PHASTER (32, 33). Antimicrobial resistance genes were detected with ResFinder 2.1 (https://cge.cbs.dtu.dk/services/ResFinder/; 34). Catabolic clusters present in the NCIMB8209 genome were searched by a BlastN approach (35) using as queries the catabolic genes previously described for the DSM300011 strain (11).

### Protein families analyses and phylogenomic calculations

The proteomes of the 126 *Acinetobacter* strains analyzed in this work, which included 99 *A. baumannii* strains other than NCIMB8209 and DSM30011, were first retrieved from the NCBI ftp database and gathered into a local database together with the NCIMB8209 proteome. Families of homologous proteins were clustered with PGAP version 1.2.1 by using the Gene Family (GF) method (36). Briefly, the BLASTALL tool was executed among all mixed protein sequences, and the filtered BLAST result was clustered by the MCL algorithm. For each protein pair of the same cluster, the global match region and the identity was set to be equal or higher than 70% of the longer sequence. This analysis allowed us to assemble 27,367 protein families among these genomes, from which 6,383 corresponded to protein families presenting a single representative in all *Acinetobacter* genomes analyzed. Regarding NCIMB8209 (3,596 predicted total CDS), 3,551 protein families contained at least one representative sequence in this strain, including 79 families that contained more than one.

Maximum-likelihood (ML) phylogenetic trees were then constructed using these data. For each protein family, the corresponding nucleotidic sequences were retrieved, individually aligned using ClustalW2 (37) and trimmed using GBlock 0.91b (38). The resulting alignments were combined to build a large supermatrix (362,285 nucleotide positions) using the Python3 package AMAS 0.98 (39). Finally, the evolutionary history of the strains was inferred as implemented in Espariz *et al*. (40) using the RAxML software (41) and the GTR substitution model with Gamma distribution. The individual parameters for the model were estimated and optimized for each concatenated gene as indicated by Stamatakis *et al*. (41). Reliability of the inferred tree was tested by bootstrapping with 100 repetitions.

In order to depurate the list of protein families specific for strain NCIMB8209 according to PGAP predictions, their coding genes were used as queries in a BLASTN sequence similarity-based comparison (35) against all 126 *Acinetobacter* spp. genomes present in our local database. Coverage and identity thresholds of 70% and an e value cut-off of 1 e^-30^ were used for these calculations. Those CDS for which a significant hit was found were then deleted from the list.

### Sequence typing

Assignment of sequence types (ST) for NCIMB8209 was done using the housekeeping genes *cpn60, gltA, gpi, gyrB, recA*, and *rpoD* (Oxford scheme) (42), or *cpn60, fusA, gltA, pyrG, recA, rplB*, and *rpoD* (Pasteur scheme) (3). For details see the *A. baumannii* MLST Databases website (http://pubmlst.org/abaumannii/) (43).

### *gdhB* gene detection by PCR

PCR reactions were carried out following standard protocols, using the primer pair GHDB_1F (GCTACTTTTATGCAACAGAGC)/GHDB775R (GTTGAGTTGGCGTATGTTGTGC) for *ghdB* detection, and the degenerate primer pair 16S RDNAF (AGAGTTTGATCHTGGYTYAGA)/16S_RDNAR /ACGGYTACCTTGTTACGACTTC) for 16S rRNA gene detection.

### Antimicrobial susceptibility testing

The general antimicrobial susceptibility of strain NCIMB8209 was evaluated using the VITEK 2 System (BioMérieux) following criteria recommended by the CLSI (Clinical and Laboratory Standards Institute, 2016, Performance standards for antimicrobial susceptibility testing. Document M100S, 26^th^ ed., CLSI, Wayne, PA). Susceptibility tests to tetracycline, chloramphenicol, and macrolides such as azithromycin and erythromycin were done separately by disk assays on Mueller-Hinton agar (MHA) following CLSI protocols. In short, NCIMB8209 cells were grown overnight at 37 °C, resuspended in LB broth to a turbidity of 0.5 McFarland units, and spread on the surface of MHA-containing Petri plates. Antibiotic disks were then carefully deposited at the center of the agar surface, and the plates were incubated at 37 °C for 16 h before measuring the diameter of the corresponding growth inhibition zones.

### Bacterial competition assays

The different *A. baumannii* and *E. coli* strains used in competition assays were grown overnight in 2 mL of L-broth (10 g/l tryptone, 5 g/l yeast extract and 0.5 g/l NaCl) medium. Cultures were then diluted in L-broth medium to Abs600 of 0.1. The cells were mixed in a 10:1 ratio (predator:prey). Aliquots of 20 μl of these cell mixtures were laid on the surface of L-Broth medium supplemented with 1.5% agar and incubated at 37°C for 4 h. The bacterial spot on the agar surface was subsequently removed and vigorously resuspended in an Eppendorf tube with 500 μl of PBS. The mixtures were serially diluted 1:10 in PBS using a 96-well polystyrene microtitre plate, and then plated onto solid selective media for colony counting. For *E. coli* DH5α, selective medium containing 20 μg/ml nalidixic acid was used. For *Acinetobacter* strains, selective media contained either 30 μg/ml gentamicin or 75 μg/ml rifampycin were employed as indicated. For all bacterial competition experiments one representative image of the growth results obtained in selective medium is shown.

### Hcp secretion analysis

Experiments were performed as previously described in Repizo *et al*. (44). Briefly, the *A. baumannii* strains analyzed for Hcp secretion were inoculated in 50 ml L-broth at an Abs600 of 0.08, and incubated at 37°C until the Abs600 reached a value of ∼1.0. Culture supernatants were obtained by centrifugation at 4,000 x g followed by filtration using 0.22 μm filters. Secreted proteins present in the supernatants were then concentrated 50-fold using Amicon Ultracel 3K centrifuge filters following the instructions of the manufacturer, and subjected to a wash step with 40 mM Tris pH 8, 200 mM NaCl, 5% glycerol before being analyzed by 18% SDS-PAGE.

### Biofilm assays

Biofilm formation was qualitatively determined by measuring the adhesion of bacteria to the surface of glass tubes. *A. baumannii* strains were statically grown overnight in 2 mL of L-Broth medium. The spent liquid medium was discarded, and the tubes were rinsed twice with water before adding a solution of 1 % Crystal Violet. After 15 min the dye solution was discarded and the tubes were rinsed twice with water before inspection. All assays were done at least three times using fresh samples each time.

### *A. baumannii* virulence assays using *Caenorhabditis elegans* nematode and *Galleria mellonella* larvae model systems

Virulence assays of the different *A. baumannii* strains tested here were evaluated by using two different model systems: *G. mellonella* moth larvae (45) and the nematode *C. elegans* (46). *G. mellonella* larvae were purchased from Knutson’s Live Bait (Brooklyn, MI) and were used the day after arrival. Groups of twenty randomly picked larvae were used for each assay condition. The different *A. baumannii* strains tested were grown overnight in LB and then diluted with PBS to obtain the CFU titers indicated in the corresponding figure legends, which were verified by colony counts on LBA for all inocula. A Hamilton microliter syringe was used to inject 10 μl of the bacterial suspensions into the hemolymph of each larva via the second last left proleg. As a control, one group of *G. mellonella* larvae was injected with 10 μl of PBS. After injection, the larvae were incubated in plastic plates at 37°C and the numbers of dead individuals were scored regularly.

For *C. elegans* survival experiments the N2 Bristol (wild type) was used. Gravid hermaphrodites were bleached and resulting eggs washed using standard protocols (47). Nematodes arrested in stage 1 (L1) were then transferred onto NGM plates seeded with fresh *E. coli* OP50 and grown at 20°C until nematodes reached larval stage 4 (L4). For virulence assays, *A. baumannii* and *E. coli* OP50 (control) were grown overnight at 37°C in L-Broth medium.

The stationary culture was then diluted to 10^7^ total CFU and 50 µl portions were seeded onto 60-mm nematode growth medium (NGM) plates. Groups of L4 worms were then transferred to the plates seeded with *A. baumannii* and deposited on the center of the bacterial spot. Assays were performed in duplicates. The nematodes were maintained at 20°C, transferred to fresh plates every 48 hours and daily monitored. Worms that did not respond to stimulation by touch were scored as dead.

Survival curves were plotted using PRISM software, and comparisons in survival were calculated using the log-rank Mantel-Cox test and Gehan- Breslow-Wilcoxon test.

## RESULTS

### Phylogenetic analysis assigned strain NCIMB8209 to *A. baumannii*, albeit to a separate clonal lineage as compared to its companion strain DSM30011

#### NCIMB8209 origins

As described in our previous work (11), strain NCIMB8209 and its companion DSM30011 were isolated prior to 1944 from the natural microbiota enriched during the aerobic decomposition of guayule, an industrial procedure designed as retting used to reduce the resinous content of the shrub processed material for the subsequent production of natural latex (14, 15). Our ML phylogenetic analyses based on core gene sequence comparisons derived from the WGS data (see below) allowed us to confidently assign this strain to *A. baumannii* as a species. In concordance with this assignment, this strain was capable of growing at 44 °C (11) in what represents a typical phenotype associated to *A. baumannii* (1, 48). Still, random amplification PCR (11), phylogenetic, comparative genome analysis, and metabolic studies (see below) indicated significant differences between this strain and its companion *A. baumannii* strain DSM30011. This evidence suggests that, even when these two *A. baumannii* strains might share similar environmental niche, they still belong to separate clonal lineages. NCIMB8209 thus provided us with a different *A. baumannii* strain isolated from a non-clinical source (14) before the massive introduction of antibiotics to treat infections (2).

#### NCIMB8209 genomic features

Genome sequencing indicated that the NCIMB8209 chromosome consisted of 3,751,581 bp in length with a G+C content of 39.1% (Table 1). These values match the average values reported for the genomes of the species composing the *Acinetobacter* genus (3,870 kpb and 39.6% respectively (6). Noteworthy, 223 of the CDS (i. e. around 6% of the total) predicted in the NCIMB8209 genome were pseudogenes. It is worth noting that a similarly high number of non-functional genes (272; 9% of the total genes) has been reported for the *A. baumannii* strain SDF, which was isolated from a human body louse and whose genome is riddled with numerous prophages and ISs (49).

**Table 1.** General features obtained from the NCIMB8209 genomic sequencing.

| **Genomic parameters<sup>a</sup>** |  |
| --- | --- |
| **Chromosome** |  |
| Accession number | CP028138 |
| Estimated size (bp) | 3,751,581 |
| Average GC content (%) | 39.1 |
| Number of CDSs | 3,596 |
| Number of tRNA genes | 73 |
| Number of rRNA genes | 18 |
| Number of ncRNA genes | 4 |
| Phage regions | 5 |
| IS copies | 79 |
| **Plasmid** |  |
| Accession number | CP028139 |
| Estimated size (bp) | 133,709 |
| Average GC content (%) | 40.1 |
| Number of CDSs | 156 |
| Number of tRNA genes | 1 |
| Number of rRNA genes | - |
| Number of ncRNA genes | - |
| Phage regions | 2 |
| IS copies | 14 |

<sup>a</sup> Based on NCBI Prokaryotic genomic annotation pipeline.

Our analysis also showed the presence of a large plasmid of 133,709 bp with a G+C content of 40.1%, hereafter designated pAbNCIMB8209_134 (Table 1). Comparison of pAbNCIMB8209_134 with other plasmids deposited in databases indicated extensive sequence identity with a group of *A. baumannii* plasmids higher than 100 kb in length, including pABTJ2 (50). All of these plasmids share a previously undescribed Rep-3 superfamily (pfam0151) replication initiation protein gene (C4×49_18465). The presence of both partition (C4×49_18550) and toxin/antitoxin genes (C4×49_18715-C4×49_18720) related to plasmid stability were detected in pAbNCIMB8209_134. On the contrary, genes involved in mobilization, conjugation, antimicrobial resistance or virulence functions could not be identified in pAbNCIMB8209_134. Still, this plasmid encodes functions which may provide some adaptive advantages to their *Acinetobacter* hosts, such as a putative glutathione-dependent pathway of formaldehyde detoxification (C4×49_18640-18650).

#### Comparisons of the chromosomal architectures of the A. baumannii environmental strains NCIMB8209 and DSM30011

Comparison of the overall chromosome structures of the *A. baumannii* NCIMB8209 and DSM30011 strains showed that the size of the former is 198 kpb smaller than that of DSM30011 (Fig. 1). Furthermore, a number of GIs and prophages distinguished these two chromosomes as will be described in greater detail below. Despite these differences a general shared synteny was observed between the two chromosomes, with the notable exception of an inversion of a 53.1 kb-region located between the first and sixth rRNA operons (Fig. 1). A similar situation was found in the community-acquired strain *A. baumannii* D1279779 (51), in which the same region was inverted as compared to DSM30011 and to most other clinical strains including the type strain ATCC17978 (Fig. S1). This rearrangement was probably mediated by homologous recombination between two oppositely-oriented rRNA operons bordering this region (51). Still, and although this inversion reverses the orientation of a number of critical housekeeping genes as well as the origin of chromosomal replication (*oriC*), it has no substantial effects on the growth rate of NCIMB8209 as compared to DSM30011 in either rich or minimal medium (data not shown).

**Fig 1.**
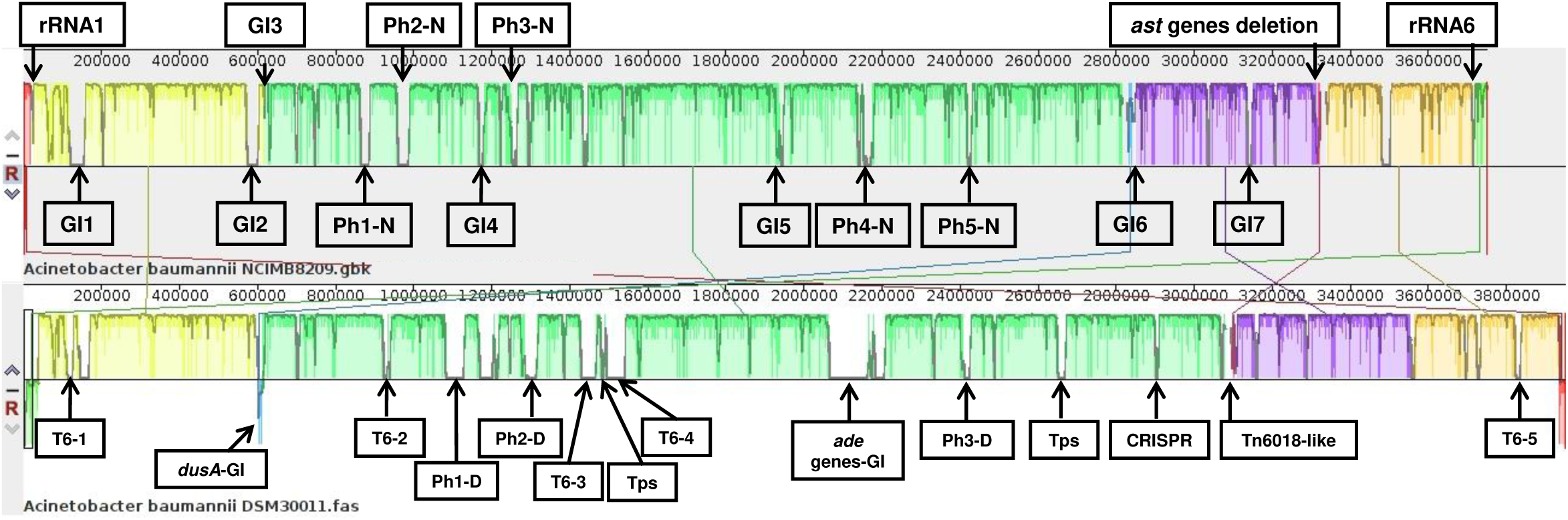
Linear comparison of the genomes of *A. baumannii* strains NCIMB8209 and DSM30011 inferred using Mauve. Each block corresponds to a DNA fragment of the chromosome distinctively coloured for clarity. The degree of conservation is indicated by the vertical bars inside the blocks. Their position relative to the genome line denotes colinear and inverted regions. For a better appreciation of genomes synteny, DSM30011 chromosomal DNA shown corresponds to the reverse complementary strand. Putative prophage (PhX-N, for NCIMB8209 and PhX-D, for DSM30011) and genomic islands (GI) insertions sites are indicated (see Tables 2 and 3 for details). Two out of the six rRNA-encoding operons in strain NCIMB8209 are also represented (rRNA1 and rRNA6). Regions encoding interbacterial competition islands (T6-1 to 5, and Tps), heavy metals (*dusA*-GI and Tn6018-like) and antimicrobial (*ade*-GI) resistance islands and a CRISPR-cas cluster are indicated for DSM30011 genome

#### Phylogenomic and MLST analyses

A phylogenetic study based on the comparisons of the concatenated sequences of 383 core genes of the environmental strains NCIMB8209 and DSM30011 and a number of *Acinetobacter* genomes that encompassed other 99 *A. baumannii* as well as 26 non-*A. baumannii* representatives (26 strains; Table S1) reinforced the affiliation of NCIMB8209 to *A. baumannii* as a species (Fig. 2 and Fig. S2). Different authors have noted the lack of a defined phylogenetic structure for the general *A. baumannii* population on phylogenetic trees based on core genes comparisons, with the exception of different terminal clusters each corresponding to an epidemic CC (2, 3, 11, 52). The incorporation of NCIMB8209 core genome sequences to this phylogenetic study did not change this general picture (Fig. 2), but some observations derived from this analysis are worth remarking. First, strains NCIMB8209 and DSM30011, which were isolated more than 70 years ago, neither emerged close to the root nor forming a separate “environmental” cluster in the *A. baumannii* subtree. On the contrary, they appeared intermixed between more contemporary clinical strains (Fig. 2). Second, NCIMB8209 emerged in a compact cluster (100 % bootstrap support) with a number of recently isolated *A. baumannii* strains that included PR07 (isolated in 2012 in Malaysia from the blood of a hospitalized patient, NCBI Bioproject PRJNA185400, direct submission) and ABNIH28, a MDR strain isolated in 2016 in the USA from a hospital closet drain (53).

**Table 2.** Genomic islands identified in the chromosome of strain NCIMB8209^a^.

| Region | Length (kbp) | Location | Integrase gene | Flanking genes (accession number) | Description |
| --- | --- | --- | --- | --- | --- |
| GI1 | 39.7 | 116596-156278 | C4X49_00725 | LysR-family transcriptional regulator (C4X49_00535)/tRNA-Gly-GCC (C4X49_00740) | Type III-restriction modification system. TA systems (HypBA, YefM-YoeB). ISs |
| GI2 | 30.5 | 569359-599853 | C4X49_02720 | hypothetical protein (C4X49_02715)/peptide chain release factor 3 (C4X49_02900) | Phagic proteins. Type I-restriction modification system |
| GI3 | 19.1 | 613054-632163 | C4X49_02950 | *dusA* (C4X49_02945)/MFS transporter (C4X49_03050) | Arsenic and heavy metals resistance island |
| GI4 | 12.6 | 1165713-1178263 | C4X49_05820 | tRNA-Ser-TGA (C4X49_05765)/hypothetical protein (C4X49_05860) | Hypothetical proteins of unknown function |
| GI5 | 19.1 | 1927268-1946350 | C4X49_09525 | hypothetical protein (C4X49_09520) /TetR-AcrR family transcriptional regulator (C4X49_09610) | Fatty acids catabolic pathway |
| GI6 | 29.3 | 2818151-2847423 | C4X49_13865 | peptide synthetase (C4X49_13745)/tRNA-Ser-TGA (C4X49_13870) | Heavy metals resistance island |
| GI7 | 13.3 | 3133811-3147160 | C4X49_15320 | sensor histidine kinase (C4X49_15265)/tRNA-Leu-CAA (C4X49_15325) | Type I-restriction modification system |

<sup>a</sup>According to NCBI and RAST annotation

**Table 3.** Prophages inserted into the chromosome of strain NCIMB8209^a^.

| Region | Length (kbp) | Predicted Proteins | Genome Position | GC (%) | Flanking genes (accession number) |
| --- | --- | --- | --- | --- | --- |
| Ph1-N | 25.8 | 35 | 860213-886091 | 40.3 | *smpB* (C4X49_04195)/phosphopantetheine adenylyltransferase (C4X49_04375) |
| Ph2-N | 28.4 | 41 | 955316-983747 | 37.7 | *ssrA* (tmRNA;C4X49_04705)/DUF927 domain-containing protein (C4X49_04925) |
| Ph3-N | 25.2 | 29 | 1240267-1265538 | 41.5 | aspartate kinase (C4X49_06155)/alanine-tRNA ligase(C4X49_06310) |
| Ph4-N | 39.1 | 32 | 2140199-2179329 | 40.5 | MFS transporter (C4X49_10600) /hypothetical protein (C4X49_10635) |
| Ph5-N | 13.9 | 11 | 2414964-2428883 | 37.7 | *gspL* (C4X49_11920)/phosphohistidine phosphatase SixA (C4X49_12010) |

<sup>a</sup>According to PHASTER predictions

**Fig 2.**
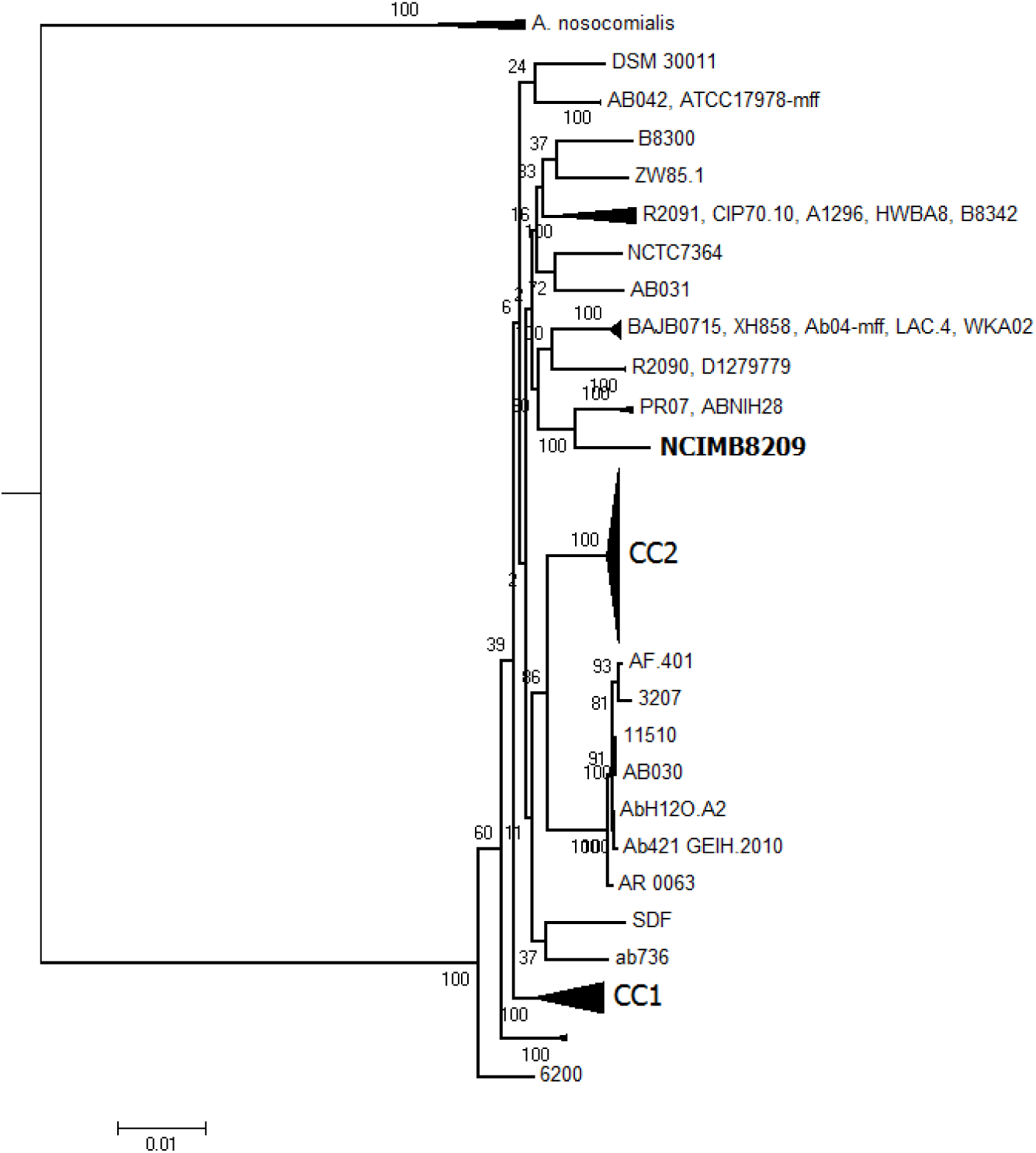
Maximum likelihood phylogenetic analysis of *A. baumannii* strains. A) The ML phylogeny was computed based on 383 concatenated core gene sequences (full tree is shown in Fig. S2). Numbers at nodes correspond to bootstrap values (100 replicates of the original dataset; BV). For simplicity, subnodes for which internal branches showed BV=100 were collapsed. The scale bar below corresponds to evolutionary distance (average number of the substitutions per site). CC1 and CC2 correspond to subclusters formed by *A. baumannii* species assigned to epidemic clonal complexes CC1 and CC2, respectively.

NCIMB8209 was assigned to a novel ST (1197, Table S1) in the Pasteur scheme of *A. baumannii* MLST classification (3). Notably, and in sharp contrast to strain DSM30011 which shared only 4 of the 7 alleles with its closest matching isolate (11), NCIMB8209 shared 6 of the 7 alleles with its closest matching isolates with all differences being restricted to the *rplB* allele (Table S1). Of note, the *gdhB* gene used in the *A. baumannii* MLST Oxford classification scheme (42) is missing from the NCIMB8209 genome (Table S1), an observation experimentally corroborated by a specific PCR assay (see Materials and Methods for details). A deeper comparative analysis with other *A. baumannii* genomes indicated that NCIMB8209 lacks a 4.8 kbp region between genes C4×49_10220 and C4×49_10225 which, in other genomes (DSM30011 included), harbors the *gdhB* gene. In the DSM30011 genome the *gdhB* gene is flanked by a hypothetical gene (DSM30011_07590) and a gene encoding a universal stress protein (DSM30011_07600), and a similar situation occurs in other *A. baumannii* genomes. This observation indicated that *gdhB* is non-essential for *A. baumannii*, and in this context its interruption by IS elements has been previously noted in a number of *A. baumannii* isolates of clinical origin (54). Moreover, a recent analysis performed with 730 *Acinetobacter* genomes revealed in 76% of them of a paralogous *gdhB* gene (*gdhB2*) in a different locus, the overall observations indicating that the Pasteur MLST scheme presents some advantages for the characterization of epidemiologically-related *A. baumannii* isolates (55).

#### NCIM8209 comparative genomics

A comparative genomic analysis that included the previously mentioned 127 different *Acinetobacter* sp. strains (Table S1) indicated that 510 representatives of the protein families detected in the NCIMB8209 genome (see Methods) are present in all of these genomes (*Acinetobacter* core genome), and 1,149 of them in all 101 *A. baumannii* genomes analyzed (*A. baumannii* core genome). These numbers differ from estimations obtained by other authors (6) of the core genome content of the *Acinetobacter* genus (950 genes; 133 genomes analyzed) and *A. baumannii* in particular (1,590 genes; 34 genomes analyzed). It is worth remarking that that even when a similar number of total genomes were used in both studies, Touchon and collaborators (2014) (6) used mostly non-*baumannii* strains whereas in our case most of the strains corresponded to *A. baumannii*. This difference and the inclusion of two *bona fide A. baumannii* environmental strains in our calculations might explain why the *A. baumannii* core genome estimation was reduced by 27.7% in our study. In this context, recent estimations carried out by other authors (56) with 78 *A. baumannii* genomes, which included the environmental strain DS002, inferred a core genome of 1,344 genes for this species. The inclusion of more environmental strains in these calculations will certainly contribute to obtain a more accurate value for the core genes repertoire of the species.

### NCIMB8209 antimicrobial resistance

Conventional antimicrobial susceptibility assays indicated that NCIMB8209 showed susceptibility to most clinically employed antimicrobials tested except nitrofurantoin and, among β-lactams, to ampicillin at MIC values just above the CLSI recommended breakpoints (Table S2). In full concordance (Table 2 and Table S2), this strain lacks AbaR resistance islands. The marginal ampicillin resistance of this strain (see above) most likely reflects the presence of a number of β-lactamase genes (Table S2). Among them, it is worth-mentioning C4×49_08235 (*bla*_OXA-78_) encoding an an OXA-51-type carbapenemase with 100 % identity to the class-D β-lactamase OXA-78 (WP_005139262.1), and C4×49_12575 (*bla*_ADC-154_) encoding an enzyme identical to the class C ADC-154 β-lactamase (WP_005138362). The presence of these genes in the chromosome of NCIMB8209 reinforces proposals that they provided for the intrinsic β-lactamase gene repertoire of *A. baumannii* (7, 57-59). NCIMB8209 also contains a *carO* gene (C4×49_13345) encoding a *carO* variant II allele described so far only among the *A. baumannii* clinical population (60), and also an *oprD/occAB1* homolog (C4×49_01130). These genes encode different outer membrane proteins proposed to participate in the permeation of carbapenems into the periplasm (61). NCIMB8209 also carries genes encoding enzymes providing resistance to aminoglycosides and chloramphenicol including *ant(3”)-II* and *catB*, respectively (Table S2). This indicates both the potentiality to evolve such resistances under selective pressure and an environmental reservoir of these resistance genes.

NCIMB8209 shares with DSM30011 (11) susceptibility to folate pathway inhibitors such as sulfamethoxazole/trimethoprim (Table S2). These susceptibilities correlate with the absence of *sul* or *dfrA* resistance genes in the genome (Table S2) and represent a notable exception when compared to most clinical strains of *A. baumannii*, including strains more contemporary to NCIMB8209 such as ATCC 19606 and ATCC 17978 (4, 7, 62-65). This characteristic is compatible with an original isolation of these two strains in an environment still free of the selection pressure derived from the use of sulfonamides common to the clinical setting at that time (11).

Genes encoding components of the RND, DMT, MATE, MFS, and SMR efflux systems involved in clinical *Acinetobacter* strains in the extrusion of toxic compounds including some antimicrobials (7) were also found in the NCIMB8209 genome (Table S2). Remarkably, a region present in most *A. baumannii* genomes encoding phage-related proteins and the AdeRS two-component regulatory system and associated AdeABC efflux components (Fig. 2) was missing in the NCIMB8209 genome. A genomic island (GI5) was instead found in an equivalent genomic location (Table 2). The loss of this gene cluster has also been documented in a number of other *A. baumannii* strains, irrespective of their clinical or environmental origins (10, 55).

### *A. baumannii* NCIMB8209 shows reduced virulence towards*Galleria mellonella* and *Caenorahabditis elegans*

*G. mellonella* moth larvae and the nematode *C. elegans* provide reliable models to study the virulence of numerous human pathogens, among them *Acinetobacter* genus species (45, 46). We thus decided to use these two models to evaluate the virulence of strain NCIMB8209 as compared to those of strain DSM30011 and the soil organism *Acinetobacter baylyi* ADP1 (24). In the *G. mellonella* model, DSM30011 showed a high virulence (44, 66), whereas a low-virulence capacity has been demonstrated for ADP1 (67). On the contrary, virulence in the *C. elegans* model has not been assessed before for this group of strains. As observed in Fig. 3, NCIMB8209 was less virulent than DSM30011 in either of these models, the latter environmental strain showing in particular a much higher capacity to kill *C. elegans*. On the contrary, NCIMB8209 killing capacity was close to that observed for *A. baylyi* ADP1 in either assay (Fig. 3). The observed differences in virulence between NCIMB8209 and DSM30011 indicate relevant phenotypic differences between them (see also following sections), regardless of their isolation as companion strains from a similar environmental origin following similar enrichment and culture protocols (14).

**Fig 3.**
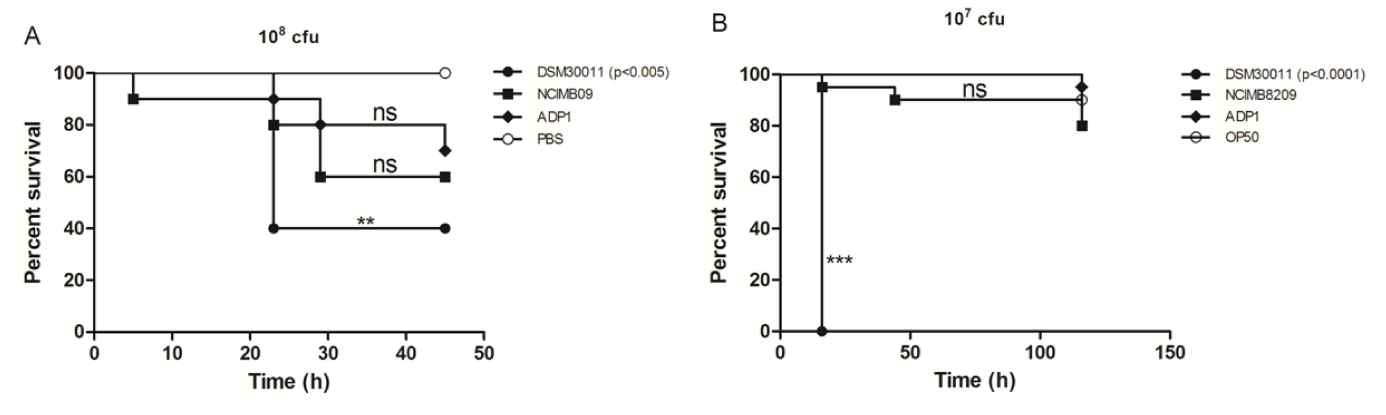
*G. mellonella* (A) and *C. elegans* (B) lethality curves. Comparative survival analysis between *A. baumannii* DSM30011 and NCIMB8209 and *A. baylyi* ADP1 strains. PBS was used as control. A) 10^8^ CFUs were used. B) 10^7^ CFUs were used. Data are representative of three separate survival experiments, each performed with 20 larvae/worms. Survival curves were constructed by the Kaplan-Meier method and compared by Log-rank analysis. n.s.: non- significant

### IS and prophages have extensively modified the NCIMB8209 genome

The lower virulence displayed by strain NCIMB8209 as compared to that of DSM30011 (see above) led us to analyze in more detail their genomes, aiming to find genetic differences that might explain these distinct phenotypes. When comparing the accessory genomes of DSM30011 and NCIMB8209 some worth-noting differences were observed in the latter (Fig. 1) such as: 1) the presence of 7 GIs (Table 2), which will be described throughout the text; 2) the presence of 5 regions harboring putative prophages (Tables 1 and S3), that will also be described in detail below; 3) the notorious absence of a CRISPRs-*cas* gene cluster common to DSM30011 and other *A. baumannii* strains (11, 68); 4) the lack of interbacterial competition islands (ICI), such as those encoding the type 6 secretion system components and/or its associated toxins (44); 5) significant differences regarding ISs number and composition between these two strains (see below).

#### Prophage content

Recent bioinformatic analyses uncovered a wide distribution and diversity of prophage contents in *A. baumannii* (69), some of them contributing to horizontal gene transfer between strains by generalized transduction (70). Our bioinformatic analysis of phage sequences in the NCIMB8209 genome indicated 5 regions (Ph1-N to Ph5-N) encompassing phage-related genes (Table 3). According to PHASTER integrity predictions, 2 “questionable” (Ph1-N and Ph2-N) and 3 “incomplete” prophages (Ph3-5-N; Table S3) were identified among them. The integration sites for Ph4-N and Ph5-N in the NCIMB8209 genome were the same hot spots found in other *A. baumannii* genomes (10), whereas those of Ph1-N to Ph3-N represent novel integration sites (Table 3). Of note, both Ph1-N and Ph5-N sequences share significant identity with the podoviral lytic bacteriophage YMC/09/02/B1251 ABA BP (NC_019541.1; (71)), while Ph2-N, Ph3-N and Ph4-N showed homology with the *Acinetobacter* phage vB_AbasS_TRS1 (NC_031098; (72)). It is also worth noting that the prophages found in NCBIM8209 shared only a limited homology with those predicted for DSM30011 (11).

#### IS content

IS elements are important drivers of genome evolution in *A. baumannii* and other pathogens, modulating antibiotic resistance gene expression, mediating genome rearrangements including deletions of substantial chromosomal regions, promoting insertional gene inactivations, etc. (73). Notably, and in sharp contrast to *A. baumannii* DSM300011 (11), a high representation of transposable genetic elements such as ISs or transposons was found in the genome of NCIMB8209. ISSaga predictions (31); Table S3 followed by manual examination corroborated the presence of 12 different ISs (totalizing 79 IS copies) in the NCIMB8209 chromosome and 9 different ISs (totalizing 14 IS copies) in the plasmid that this strain harbors (Table 4). These values are significantly higher than the estimated average of 33 IS copies per *A. baumannii* genome determined by Adams and collaborators (8). Moreover, it is also remarkable the diversity of IS families (16 in total) found in the NCIMB8209 genome when considering that only 7 out of 976 *A. baumannii* genomes were found to carry 10 or more different IS elements (8). Although most of the IS elements identified in NCIMB8209 were previously reported in different species of the *Acinetobacter* genus, this strain contains a very particular IS profile for an *A. baumannii* strain, with 2 of the most frequent IS elements (ISAha2 and ISAha3) originally detected in *A. haemolyticus* (Table 4). Furthermore, as also seen in this Table, NCIMB8209 harbors 2 of the 5 most represented ISs in *A. baumannii* (8), namely ISAba26 (15 copies) and ISAba13 (14 copies). It is also remarkable the absence of ISAba1 copies in NCIMB8209, considering the large impact of this IS among clinical *A. baumannii* genomes (8). Of note, the 3 most represented chromosomal ISs (ISAha3, ISAba26 and ISAba13) are also carried by plasmid pAbNCIMB8209, suggesting that this plasmid was the vehicle enabling their acquisition.

**Table 4.** ISs content^a^.

| Denomination | Origin | DNA molecule |  |
| --- | --- | --- | --- |
|  |  | Chromosome<sup>b</sup> | pAbNCIMB8209 |
| ISAha3 | *A. haemolyticus* | 18 (2) | 1 |
| ISAb26 | *A. baumannii* | 15 (4) | 2 |
| ISAb13 | *A. baumannii* | 14 (7) | 3 |
| ISAb44 | *A. baumannii* | 11 (2) | - |
| ISAha2 | *A. haemolyticus* | 7 (2) | - |
| IS1006 | *A. junnei* | 4 | - |
| ISAb45 | *A. baumannii* | 3 | - |
| IS17 | *A. haemolyticus* | 2 (1) | - |
| IS1008 | *A. calcoaceticus* | 2 (1) | - |
| ISAb22 | *A. baumannii* | 1 (1) | 2 |
| ISAb4 | *A. baumannii* | 1 | - |
| ISAb31 | *A. baumannii* | 1 | 2 |
| ISAb2 | *A. baumannii* | - | 1 |
| ISAb46 | *A. baumannii* | - | 1 |
| IS1007 | *Acinetobacter sp.* | - | 1 |
| ISAcsp3 | *Acinetobacter sp.* | - | 1 |

<sup>a</sup> According to ISSaga and ISFinder predictions
<sup>b</sup> Number of gene-interrupting IS copies are indicated in parenthesis

From the detected ISs in the NCIMB8209 genome, 3 represented novel elements which were deposited at the ISSaga database (31) under the names ISAba44 (IS630 family), ISAba45 (IS3 family), and ISAba46 (IS66 family), respectively. From them, ISAba44 is present in 11 copies in the NCIMB8209 chromosome (Table 4). This particular mobile element was also found in the chromosome of other 3 *A. baumannii* strains (ABNIH28, B8300 and B8342) and in *A. johnsonnii* XBB1 (75% identity, query coverage=100%). No homologs to this IS were found in other organisms outside the *Acinetobacter* genus. ISAba45 (3 copies) was also detected in other four *A. baumannii* strains (PR07, ABNIH28, A1296, and IOMTU433, the former two strains closely linked to NCIMB8209, Figure 2) and in *A. soli* GFJ2 (87% identity, query coverage=99%). Moreover, ISAba45 displays significant nucleotide identity with similar ISs present in other *Acinetobacter* strains (76%) and in *Moraxella osloensis* plasmids (70%). The ISAba46-only copy is carried by plasmid pAbNCIMB8209, and was also found in the chromosomes and plasmids of numerous strains of *A. pittii, A. baumannii, A. lwoffii, A. junii, A. johnsonnii* and *A. haemolyticus* (>86% identity, query coverage=100%). Concerning the locations of the above IS elements, we found that 17 of them are interrupting an equivalent number of CDSs, very likely precluding their expression (Table S3. Seven of these 17 CDSs are interrupted by ISAba13 copies (Table 4). Remarkably, a similar situation was reported previously for strain D1279779, which harbors 18 ISAba13 copies (51). We also detected that some IS elements were positioned on one of the borders of several GIs, including GI1, GI2, GI4, and GI6 (Table 2). It is tempting to speculate that these insertions probably interfere with the excision mechanism of the corresponding GIs, and were thus selected due to the subsequent retention of these GI in the chromosome (74).

In summary, the plethora of IS elements found in NCIMB8209 have significantly help remodeling the genome of this strain.

#### A genome devoid of interbacterial competition islands

As mentioned above, the genome of NCIMB8209 neither carries a T6SS-main gene cluster nor any T6SS-associated locus encoding VgrG-like proteins or their cognate toxins (44, 75). Searching for other competition mechanisms such as the Two-partner systems (Tps) already identified in strain DSM300011 (Fig. 2) related to contact-dependent inhibition (CDI) (76), also resulted in negative results. These observations led us hypothesize that the NCIMB8209 ability to outcompete other bacteria was severely compromised. To test this prediction, bacterial competitions assays were performed essentially following previously described procedures (44). Results shown in Fig. S3 demonstrate that NCIMB8209 was not capable of outcompeting *E. coli* when the former strain was used as the attacker and *E. coli* DH5α was the prey. On the contrary, the DMS30011 strain was clearly capable of outcompeting *E. coli* DH5α in a similar assay, an effect that specifically depended on the T6SS system as shown by the lack of effect observed in a DMS30011 *ΔtssM* mutant (Fig. S3A). In correlation with these results NCIMB8209, similarly to DSM30011 *ΔtssM*, did not secrete detectable amounts of Hcp (a marker of a functional T6SS) into the growth medium (Fig. S3B). We additionally tested the capacity of NCIMB8209 and DSM30011 to out-compete each other. While DSM30011 completely eliminated NCIMB8209 (Fig. S3C) when co-incubated in a 10:1 ratio attacker:prey, the latter strain was not capable of outcompeting the former under similar experimental conditions (Fig. S3D).

#### Persistence and virulence

To analyze the presence of genes potentially involved in persistence and virulence in strain NCIMB8209, we performed a BlastN-homology search using as query a list of potential candidates described in *Acinetobacter* strains (11). This list included genes coding for the synthesis of the capsule and other exopolysaccharides, appendages, OM proteins and the T2SS; genes coding phospolipases and proteases; and genes involved in traits such as motility and iron scavenging. Our searching indicated that 131 out of the 146 genes (sequence identity ≥76%) encoding potential virulence factors analyzed were present in NCIMB8209 (Table S2). Among the 15 putative virulence factors included in the search and not detected in the NCIMB8209 genome, most of them encode or are involved in the synthesis of surface-exposed molecules (Table S2). These included the Prp pilus (77), the MFS transporter Pmt (probably involved in DNA transport necessary for biofilm formation; (78)) and a surface motility-associated molecule (79). Moreover, the gene encoding for the Pilus 3-fimbrial adhesion precursor (C4×49_08175) and the Pilus 3-fimbriae anchoring protein (C4×49_08180) are both incomplete, and a gene coding for a polymorphic toxin of the RTX-family (80) was interrupted by an ISAba44 copy (C4×49_17180-C4×49_17185). We also noticed that several genes which products are directly or indirectly involved in functions related to adhesion or biofilm formation in clinical strains were absent or interrupted by ISs. For instance, the gene encoding the Bap protein (81) could not be detected (Table S2), and instead an ISAba13 copy was identified in the locus encompassing the *bap* gene in other *A. baumannii* strains. Other examples are the gene coding for the Ata adhesin (82) which was interrupted by an ISAba26 copy, and a gene encoding the O-antigen ligase TfpO normally involved in glycan decoration of capsule and proteins, which was interrupted by an ISAba13 copy in NCIMB8209 (83, 84). We then hypothesized that the lack of this group of genes could have impacted traits such as biofilm and pellicle formation as well as motility. To evaluate this hypothesis, the capacity of NCIMB8209 to form pellicle and biofilm when grown overnight in rich medium was compared to that of DSM30011, which was found to represent a high-biofilm/biopellicle producer (44). This assay indicated that NCIMB8209 was neither capable of forming pellicle (Fig. S4A) nor attaching to glass surfaces (Fig. S4B), supporting the above prediction that the ability to form biofilm/biopellicles is severely compromised in this strain. Furthermore, motility assays on semisolid medium showed that the ability of NCIMB8209 to perform swarming was highly reduced (Fig. S4B), again in sharp contrast to DSM30011 (66).

Another gene absent in NCIMB8209 is *cpaA* (Table S2), which encodes the surface-exposed metallopeptidase Cpa endowed with the ability to cleave fibrinogen and coagulation factor XII, and thus proposed to deregulate blood coagulation (85, 86). CpaA represents a substrate of the Type II secretion system (87), and has been proposed as a *bona fide A. baumannii* virulence factor (88). Therefore, its conspicuous absence in NCIMB8209 (Table S2) and also in its companion DSM30011, as well as in the clinical strains ATCC 17978 and ATCC 19606 isolated by the middle of last century (11), supports previous proposals that this gene was recently acquired by *A. baumannii* by horizontal gene transfer (87).

### Genetic features contributing to NCIMB8209 environmental adaptation

#### Gene acquisition by lateral gene transfer

Our genomic comparative analysis and subsequent BlastN-based homology search (see Methods) indicated that 77 CDSs (59 of them located in the chromosome and the extra 18 in the plasmid) are unique to NCIMB8209 (Table S4). From the 59 unique chromosomal genes, 38 were located within different prophage and GI regions and showed no significant hits in databases. A Blast searching against the NCBI Protein database using as query the amino acid sequence of the remaining 21 chromosomal CDSs revealed that 20 of them had significant best hits with proteins found in the database, sixteen of them (i. e., 80%) located in species of the *Acinetobacter* genus (12 of them in *A. baumannii*) with potential roles in transport, motility, transcriptional regulation, and lipopolysaccharide synthesis (see Table S4 for details). Notably, 4 CDSs related to capsule synthesis (see below) were best affiliated to homologs located either in species of different orders among the class *Gammaproteobacteria* to which *Acinetobacter* belongs, including the *Alteromonodales* (*Alteromonas sp*.) and the *Enterobacteriales* (*Yersinia sp*.), and also to a different class (*Azoarcus sp*., *Betaproteobacteria*) or even to a different phylum (*Vitellibacter sp*., *Bacteroidetes/Chlorobi*) (Table S4). This indicated a remarkable ability of *A. baumannii* NCIMB8209 to co-opt genes from both phylogenetically-related and - distant species as the result of horizontal gene transfer.

#### NCIMB8209 carries a novel K-locus

Among idiosyncratic features of NCIMB8209 worth remarking, we found differences in content and organization of the genes linked to the production of the K capsule. The K locus identified in the NCIMB8209 genome (Table S2) displayed a gene organization that resembles the polysaccharide gene cluster PSgc6 reported for other clinical *A. baumannii* strains (89), with the exception of the *wafQRST* cluster which is missing in NCIMB8209. Furthermore, three extra genes were found in this novel K locus, namely C4×49_00290 encoding an O-acetylase (absent in other members of the *Acinetobacter* genus), C4×49_00300 encoding a glycosyl-transferase, and C4×49_00305 encoding a pyruvyl transferase (Table S2). Pyruvyl-capped N-acetyl-D-galactosamine (D-GalpNAcA) branches constitute rare structures described so far only in *A. baumannii* D78, a strain assigned to CC1 (90). The K-locus arrangement described above for strain NCIMB8209 was not found in other *Acinetobacter* strains by database searching, therefore revealing a previously unreported PSgc locus in *A. baumannii*. Of note, the *weeK* gene coding for an UDP-N-acetyl-glucosamine 4,6-dehydratase (involved in the biosynthesis of UDP-linked sugar precursors used for capsule synthesis) is also interrupted by an ISAba44 copy in NCIMB8209 (C4×49_00335-C4×49_00350, Table S2).

NCIMB8209 shares with DSM30011 a similar gene locus involved in the synthesis of outer core polysaccharides (OC) of the lipid A core moiety, with the exception of an additional gene encoding a glycosyl transferase (C4×49_15205). However, this gene is annotated as a pseudogene and is probably non-functional (Table S2). As previously described for strain DSM30011 (11), this cluster includes the *rmlBDAC* genes (Table S2) responsible of the biosynthesis of dTDP-L-rhamnose (89, 91). The observed genetic organization of the OC locus in NCIMB8209 corresponds to OCL6 according to the classification proposed by Kenyon and collaborators (92).

#### Mechanisms of resistance to toxic compounds

NCIMB8209 also contains many gene clusters encoding systems involved in the resistance to toxic compounds. Some of these clusters are scattered throughout the genome, while others are concentrated in three regions. One of these regions is constituted by GI3 (Table 2 integrated next to the *dusA* gene (C4×49_02945). The *dusA* locus has been found to represent a common integration site for this kind of genetic islands in *A. baumannii* (93, 94). Although GI3 (19.1 kb long) is shorter than homologous GIs carried by other *A. baumannii* strains (for instance, in DSM30011 this GI is 33 kb long), this island still includes genes for putative arsenate and heavy metal ion detoxification systems (*ars* and *czc* genes) and other involved in Fe ions transport (*feoAB*). Another case is GI6, which carries a cluster of genes encoding a putative copper ion detoxification system (C4×49_13800-C4×49_13840, Table 2). More interestingly, GI6 also harbors a *mobA* gene (C4×49_13855) encoding a protein with a relaxase domain which might be responsible for its mobilization after excision from the genome (Table 2). This may suggest a plasmid origin for this GI, and also opens the possibility that this gene could even mediate its mobilization by horizontal gene transfer after excision from the genome. The third cluster (C4×49_16225-C4×49_16400) contains a *merR-merTPCAD* gene cluster (C4×49_16305-C4×49_16300 to C4×49_16280) coding for a complete Hg ion detoxification system (95). Since there is no gene coding for an integrase nearby, this region was not considered as a GI. Remarkably however, it is flanked by several IS copies which might have contributed to its mobilization and integration in this locus.

Inspection of the NCIMB8209 genome also evidenced the presence of 3 putative catalase genes (C4×49_07525, C4×49_02120 and C4×49_17825). Catalases represent one of the main strategies evolved by cells to cope with the accumulation of reactive oxygen species (26). It is then noteworthy that the number of putative catalase proteins encoded by this strain is higher than that of the environmental strains DSM30011 (2, PNH13446.1 and PNH14300.1) and *A. baylyi* ADP1 (1, CAG67388.1), and similar to that found in the *A. baumannii* clinical strain ATCC 17978 (3, ABO11814.2, ABO10867.2 and ABO13771.2). Moreover, a comparative analysis of the tolerance of the above strains to strong oxidants, as judged by their survival when exposed to H_2_O_2_ (26), indicated a strong correlation between their catalase gene content and oxidative stress resistances (Fig. S6).

#### Catabolic abilities

Previous WGS analysis of the environmental *A. baumannii* DSM30011 strain predicted the presence in its genome of 28 gene clusters encoding many metabolic pathways involved in the utilization of a large variety of plant substances (11). The presence and organization of similar catabolic genes was also investigated in NCIMB8209 and, despite some differences in the organization of catabolic loci between these two strains, 27 out of the 28 catabolic loci found in DSM30011 (11) were also present in this strain. The only exception was the salicylate/gentisate (*sal2*/*gen*) cluster, which was totally missing in NCIMB8209. In addition, the *betABI* locus present in both strains (previously thought as a catabolic gene cluster, (11, 24)) has been shown in *A. baylyi* to be involved in the synthesis rather than in the degradation of glycine betaine (96). In agreement, none of the *A. baumannii* strains tested including DSM30011, NCIMB8209, and ATCC 17978, nor *A. baylyi*, were capable of utilizing glycine betaine as the only carbon source for growth (Table S5).

From the 27 predicted catabolic *loci* shared between NCIMB8209 and DSM30011 mentioned above, 17 equivalent clusters are also found in the genome of *A. baylyi* and are involved in the degradation of plant substances and the recycling of plant material (24). These include loci such as *pca, qui, pob, hca, van*, and *ben*, involved in the degradation of aromatic acids and hydroxylated aromatic acids such as hydroxycinnamic acids constituting the building blocks of plant protective heteropolymers such as suberin (24, 97-99). These aromatic compounds are ultimately catabolized through the beta-ketoadipate pathway yielding Krebs cycle-intermediary substrates, therefore allowing bacterial growth when used as substrates (24). Our analysis of the ability of *A. baumannii* strains NCMIB8209 and DSM30011 to employ different compounds as substrates for growth (Table S5) indicated that these two strains share with *A. baylyi* the ability to utilize many aromatic acids found in plants including benzoate, 4- hydroxy-benzoate, 4-hydroxy-cinnamate, and shikimate, as sole carbon sources. Moreover, the activity of the *mdc* pathway involved in the catabolism of dicarboxylic malonic acid, another plant-synthesized compound (24), was inferred from the growth observed by all *A. baumannii* strains tested in malonate as the only carbon source (Table S5). These observations are compatible with the isolation of *A. baumannii* strains strains NCMIB8209 and DSM30011 from an enriched consortium specialized in the recycling resinous plants material (14). Remarkably still, all of the above-mentioned catabolic capabilities are also shared by *A. baumannii* clinical strains such as ATCC 17978 (Table S5 and data not shown).

Besides the above described similarities with *A. baylyi*, NCMIB8209 and DSM30011 are endowed with some idiosyncratic catabolic clusters also related to the degradation of particular plant compounds. Among them we could mention the *paa* (phenylacetic acid, PAA) and *liu* (leucine/isovalerate) clusters (11). The presence of a *paa* cluster in both NCMIB8209 and DSM30011, but not in *A. baylyi*, correlates with the capability of these *A. baumannii* strains to grow on PAA as the sole carbon source (Table S5). PAA is a plant auxin derived from the catabolism of phenylalanine (100, 101) endowed with substantial antimicrobial activity (102). PAA degradation by *A. baumannii* clinical strains has already been noted (100) (see also Table S5) and found to play an important role during *A. baumannii* infection by reducing the levels of this powerful phagocyte chemoattractant (101). Concerning the *liu* (leucine/isovalerate) catabolic cluster, evidence of its activity was obtained by the growth observed for NCMIB8209 and DSM30011 on L-leucine or isovalerate as only carbon sources (Table S5). In *P. aeruginosa* the *liu* pathway complements the *atu* pathway responsible of the degradation of acyclic terpenes produced by plants in response to phytopathogens (103), and a similar situation may occur also in NCMIB8209 and DSM30011 (11). It follows that these *A. baumannii* strains have the capacity not only to participate in the degradation of many plant aromatic compounds, but are also endowed with the additional ability to degrade compounds produced by plants in response to stress situations including the attack of phytopathogens and phytophagous insects (97-99).

Besides the above described similarities at both the genomic level and metabolic capabilities between DSM30011 and NCIMB8209, a differential capacity for the utilization of the basic amino acids arginine and ornithine was observed between these two strains (Table S5). This may be explained by the differential presence in DSM30011 of a 10.5 kbp fragment containing a *gdh/ascC/astA/astD/astB/astE* gene cluster encoding a complete arginine succinyltransferase (AST) pathway responsible of the catabolism of these basic amino acids (104), which was missing in NCIMB8209 (Fig. 1). Instead, a region of 27.6 kb bordered by IS1008/ISOur remnants and encompassing putative ion mercury detoxification genes was found in NCMIB8209, which has in turn been heavily impacted by several ISs of different types (Table S3). It is worth noting in the above context that the chromosomal region adjacent to *astA* in *A. baumannii* CC2 strains is also an integration site for different mobile elements such as AbGRI2-type resistance islands among others, which have provoked different rearrangements in their vicinity including various deletions (10, 73). This has resulted in some CC2 strains in which a AbGRI2-type element is found adjacent to a complete *ast* gene cluster, and other strains in which the *ast* genes have been completely deleted (10, 73) (and data not shown). It has been shown in *P. aeruginosa* that the N-succinyl transferase AstA, the first enzyme of the AST pathway, is able to use both L-arginine and L-ornithine as substrates (105), and a similar substrate specificity may also occur for *A. baumannii* AstA. Furthermore, genes coding for a putrescine importer (*puuP*) and a gamma-aminobutyraldehyde dehydrogenase (*patD)*, the latter part of the transaminase pathway of putrescine degradation (104), were identified in DSM30011 (51% and 48% identity with the corresponding proteins from *Escherichia coli* str. K-12 substr. MG1655; accession numbers AAC74378.2 and AAC74526.1, respectively) but not in NCIMB8209. These latter observations might also explain the inability of NCIMB8209 to grow on putrescine, as compared to the other *A. baumannii* strains tested (Table S12). In any case, the observed phenotypic profiles suggest a narrowing of NCIMB8209 substrate utilization capabilities when basic amino acids and polyamines in particular are considered, which suggest an adaptation of this strain to a specific niche.

## CONCLUSIONS

*A. baumannii* NCIMB8209 represents to our knowledge the second reported environmental *A. baumannii* strain, isolated from a desert plant source at the onset (or even before) the massive introduction antimicrobials to treat infections (11). In concordance, and similarly to its companion strain DSM30011, NCIMB8209 showed general susceptibility to most clinically-employed antimicrobials including folate pathway inhibitors (this work and 11). Expectedly from their common plant source and retting enrichment before isolation in media containing guayule resins as substrates for growth (14), both *A. baumannii* strains share the ability to degrade a number of substances produced by plants including many hydroxylated aromatic acids constituting the building blocks of plant protective hydrophobic heteropolymers and also repellents against predators (Table S5). However, WGS and subsequent phylogenetic and comparative genome analysis, as well as different biochemical studies conducted in this work, indicated significant differences between these two environmental strains. First, as compared to DSM30011, NCIMB8209 has undergone a significant genome reduction and lacks many genetic clusters encoding components involved in defense mechanisms against other biological competitors such as the CRISPR-cas complex, T6SS, and two-partner systems. Moreover, and although NCIMB8209 contains most genes associated to persistence and virulence in *A. baumannii* clinical strains, many encoding components of surface structures are interrupted by IS elements whose relatively high number and variability impacted heavily on the NCIMB8209 genome. Among IS-interrupted genes we found those encoding pili components, the O-antigen ligase TfpO, a biosynthetic route for surfactants compounds, Bap and Ata adhesins, a RTX-toxin, etc. (Table S3). Comparative biofilm and pellicle production, as well as motility assays, evidenced in fact that NCIM8209 is severely compromised in these pathogenicity-associated traits (Fig. S4). Altogether, these observations can explain the low relative virulence potential observed for NCIM8209 on the *G. mellonella* and *C. elegans* infection models (Fig. S3).

Loss of T6SS genes has been observed in a number of *A. baumannii* clinical strains causing infections, suggesting that this system is not required once *A. baumannii* invades its host (73, 106, 107). Moreover, T6SS absence has been linked with higher chances of evasion of *A. baumannii* from the host immune system (108, 109). The situation above described for strain NCMIB8209 therefore resembles that reported for *A. baumannii* SDF isolated from a human body louse (49), which has undergone extensive genome reductions and rearrangements mediated by ISs (49). Also, albeit more moderately, loss of T6SS and IS-mediated inactivation of genes encoding surface structures was also observed in *A. baumannii* D1279779, a community acquired strain isolated from the bacteraemic infection of an indigenous Australian (51).

It is tempting to speculate that the changes observed in *A. baumannii* NCMIB8209 are also related to its adaptation to a particular niche. In this context, several reports have shown the profound associations existing between different bacterial groups including many members of the *Acinetobacter* genus with a number of insects feeding on plants (16-22). Many of these bacterial species are located in the gut of their insect hosts conducting mutualistic or symbiotic associations based on nutrition/protection relationships (16, 18-20, 110, 111). The bacterial counterpart thus degrades toxic compounds for the insect of the plant diet, and the insect host in return provides a stable environment, supply or resources, and a vector for the rapid spreading and inoculation into fresh plant tissues. For those bacterial species displaying low pathogenic potential, selective pressure eventually favours more stable relationships with concomitant structural and metabolic changes (111). A general trend thus observed for Gram-negative species with these characteristics is genome reduction with the loss or modification of surface-exposed molecules, thus reducing interactions with the innate immune system of their hosts which may trigger defensive responses and elimination (110, 111). This situation that can be certainly applied to NCIMB8209, as extensively detailed above. In this context, we note the relative high tolerance of this strain to pro-oxidants such as H_2_O_2_ (Fig. S6), which correlates with the identification of 3 catalase genes in its genome (112). The innate immune system of insects closely resembles that of vertebrates at both the molecular and cellular levels (110). Thus, while the cellular immune response in vertebrates is mediated by professional phagocytes, in insects this function is conducted by phagocytic cells known as haemocytes (110). Since phagocytic cells use oxidative burst as a common strategy to counteract pathogens (110, 113), it is then possible that the selection of a higher anti-oxidant ability in NCIMB8209 led to an increased tolerance to oxidative stress and therefore increased survival in an insect niche. Of note, recent evidence indicates that *A. baumannii* strains with high catalase production are more resistant against intracellular killing by macrophages (114).

Our observations also provide further clues on the high genomic plasticity of *A. baumannii* as a species (6), underscoring the fact that this highly advantageous feature in adaptive terms is not exclusive to *A. baumannii* strains of clinical origin only. In summary, WGS-analysis and complementary phenotypic characterization of *A. baumannii* strains of environmental origin such as DSM30011 and NCIMB8209 provided important clues on the genomic content and diversity of this species before the strong selection pressure associated to the current antibiotic era, and suggested potential niches of this species outside the clinical setting.

## AUTHOR CONTRIBUTION STATEMENT

GDR, AMV and HAS conceived and designed the work. GDR and ME conducted the bioinformatic analysis. GDR, JLS, JIDM, BAS, and AMV performed the experimental part. GDR, ME and AMV analyzed the data. All authors contributed to the writing of the manuscript, and read and approved this final version.

## ACKNOWLEDGEMENTS

This work was supported by grants awarded to AMV from the Agencia Nacional de Promoción Científica y Tecnológica (ANPCyT); Consejo Nacional de Investigaciones Científicas y Técnicas (CONICET-PIP1055); Ministerio de Ciencia, Tecnología e Innovación Productiva, Provincia de Santa Fe; Argentina; and to HAS, NIH grant AI-115203; and by a Fulbright fellowship awarded to GDR. We are indebted to the curator teams of both the Institut Pasteur MLST system (Paris, France) and Oxford University (Oxford, UK) for their help in incorporating NCIMB8209 MLST alleles and profiles at http://pubmlst.org/abaumannii/. We are indebted to D. de Mendoza (IBR-CONICET, Rosario, Argentina) for the kind gift of the *C. elegans* N2 strain. We also thank Dr. V. Müller (Goethe University Frankfurt, Frankfurt am Main, Germany) for calling our attention on the biosynthetic characteristics of the *Acinetobacter bet* locus. ME, AMV and GDR are staff members of CONICET. HS is member of the Department of Microbiology at the University of Chicago.

## List of Supplementary Tables

**Table S1:** *Acinetobacter* strains used for phylogenetic and comparative studies and MLST classification data.

**Table S2:** Genes putatively contributing to antimicrobials resistance and virulence in *A. baumannii* NCIMB8209.

**Table S3:** Prophage and IS-related genes in *A. baumannii* NCIMB8209.

**Table S4:** Unique genes carried by *A. baumannii* NCIMB8209.

**Table S5:** Catabolic tests

